# Metalation calculators for *E. coli* strain JM109 (DE3): Aerobic, anaerobic and hydrogen peroxide exposed cells cultured in LB media

**DOI:** 10.1101/2022.05.06.490408

**Authors:** Andrew W. Foster, Sophie E. Clough, Zeynep Aki, Tessa R. Young, Alison R. Clarke, Nigel J. Robinson

**Affiliations:** Department of Biosciences, Durham University, Durham, UK; Department of Chemistry, Durham University, Durham, UK; Advanced Research Computing, Durham University, Durham, UK

**Keywords:** Metal-binding, metal speciation, metal specificity, metal availability, protein expression, industrial biotechnology

## Abstract

Three web-based calculators, and three analogous spreadsheets, have been generated that predict *in vivo* metal occupancies of proteins based on known metal affinities. The calculations exploit estimates of the availabilities of the labile buffered pools of different metals inside a cell. Here, metal availabilities have been estimated for a strain of *E. coli* that is commonly used in molecular biology and biochemistry research, for example in the production of recombinant proteins. Metal availabilities have been examined for cells grown in LB medium aerobically, anaerobically and in response to H_2_O_2_ by monitoring the abundance of a selected set of metal-responsive transcripts by qPCR. The selected genes are regulated by DNA-binding metal sensors that have been thermodynamically characterised in related bacterial cells enabling gene expression to be read-out as a function of intracellular metal availabilities expressed as free energies for forming metal complexes. The calculators compare these values with the free energies for forming complexes with the protein of interest, derived from metal affinities, to estimate how effectively the protein can compete with exchangeable binding sites in the intracellular milieu. The calculators then inter-compete the different metals, limiting total occupancy of the site to a maximum stoichiometry of 1, to output percentage occupancies with each metal. In addition to making these new and conditional calculators available, an original purpose of this article was to provide a tutorial which discusses constraints of this approach and presents ways in which such calculators might be exploited in basic and applied research, and in next-generation manufacturing.

## Introduction

Metalation is difficult to comprehend based on metal affinities because proteins commonly bind wrong metals more tightly than those required for activity^1-3^. Metalation *in vivo* is sometimes assisted by metallochaperones and chelatases^4,5^, but here the question becomes, how do the correct metals somehow partition onto the delivery proteins? The order of binding of essential and exchangeable cytosolic metals to proteins typically follows the Irving Williams series^3,6^.

Cells contain vast surpluses of binding sites for most metals and a sub-set of these sites are labile^7^. Metals can transfer from labile binding sites by ligand exchange reactions^7-9^. To obtain a specific metal a protein must compete with these labile binding sites. To make metalation predictable it is necessary to know how tightly the labile metals are bound. Metal-sensors have evolved to respond to changes in metal availability and have been used to estimate how tightly the exchangeable metals are bound^10^. A set of bacterial DNA-binding metal sensors (from *Salmonella enterica* serovar *Typhimurium* strain SL1344) was thermodynamically characterised (determining *K*_metal_ for the tightest allosteric site, *K*_DNA_ of apo-sensor, *K*_DNA_ of holo-sensor, number of sensor molecules per cell in the absence and presence of elevated metal, number of promoter DNA targets) to relate DNA occupancy to intracellular metal availability. These data confirmed that the available labile pools of the tighter binding metals such as Ni^2+^, Zn^2+^ and Cu^+^ are maintained at the lowest free energies for forming metal complexes, while the weaker binding metals such as Mg^2+^, Mn^2+^ and Fe^2+^ are at the highest^10^. In short, metal availabilities in cells also follow the Irving Williams series^10^, and metalation can be understood by reference to these free energy values^10,11^.

A metalation calculator for an idealised *Escherichia coli* cell was previously created based on metal-availabilities at the mid-points of the ranges for each metal of the similar set of *Salmonella* sensors^11^. This calculator first determines the difference between these availabilities and the free energy for forming a metal-complex with a protein of interest: the latter determined from metal affinities using the standard relationship *ΔG* = -*RT*ln*K*_*A*_. The metal with the largest favourable free energy gradient, from the available labile pool to the protein, becomes the predominant metal bound. The calculator competes the free energy differences for all metals such that the total amount of metal bound to a site does not exceed a stoichiometry of one^11^.

Here we create calculators for conditional, rather than idealised, cells based on the status of the metal-sensors, and hence metal availabilities, during standard growth of *E. coli* in LB medium. *E. coli* strain JM109 (DE3) has been used for this work since it is widely exploited for molecular biology and contains a full complement of metal sensors (unlike strain BL21 for example which is aberrant in Ni^2+^ and Co^2+^ sensing and homeostasis)^12^.

Previous work calibrated the availabilities of cobalt and zinc in conditional cells of a strain of *E. coli* that had been engineered to produce vitamin B_12^11^_, and here we replicate this approach in JM109 (DE3) for all metals. Calculators have been generated for cells grown under aerobic conditions, anaerobic conditions and after exposure to H_2_O_2_. These calculators can be used to predict and optimise the metalation of recombinant proteins overexpressed in *E. coli*. They can be used to explore fundamental questions, for example related to the effects of oxygen status on metalation and to identify disparities which illuminate the contributions of more elaborate mechanisms to the specificity of metalation. A half of enzymes require metals, and it is intended that accessible calculators will assist the optimisation of metallo-enzyme dependent sustainable manufacturing in industrial biotechnology. Here we discuss constraints associated with this approach and set out ways in which outputs of the calculators might be interpreted.

## Methods

### Bacterial strain maintenance/growth and reagents

*E. coli* strain JM109 (DE3) was purchased from Promega. Liquid growth media and cultures were prepared in acid washed glassware or sterile plasticware to minimise metal contamination. Overnight cultures were inoculated into 400 ml LB (10 g/l tryptone, 5 g/l yeast extract, 10 g/l NaCl) at a 1 in 100 dilution and grown at 37 °C to an OD_600 nm_ of 0.2-0.3 then split into 5 ml aliquots in 12 ml capped culture tubes with the indicated treatment and grown as indicated. All aerobic cultures were incubated with shaking at 180 rpm while anaerobic cultures were incubated statically in an air-tight box with AnaeroGen anaerobic gas generating sachets purchased from Thermo Scientific. OD_600 nm_ measurements were made using a Thermo Scientific Multiskan GO spectrophotometer and percentage growth of n = 3 biological replicates (unless stated otherwise) calculated relative to untreated control cultures (n = 3 biological replicates).

All metal stocks (other than FeSO_4_ stock used for 180 min anaerobic exposure to 0.5 mM FeSO_4_) were quantified by ICP-MS. ICP-MS analysis was performed using Durham University Bio-ICP-MS Facility. MnCl_2_, CoCl_2_, NiSO_4_, CuSO_4_, ZnSO_4_ were prepared in ultrapure water. FeSO_4_ was prepared in 0.1 N HCl and diluted when required using ultrapure water^13^. Dimethylglyoxime (DMG) was dissolved in 100% ethanol. Metal solutions and EDTA were filter sterilised prior to addition to bacterial cultures.

### Determination of transcript abundance

Aliquots (1-1.2 ml) of culture were added to RNAProtect Bacteria Reagent (Qiagen) (2-2.4 ml), vortexed and incubated at room temperature for 5 min before pelleting by centrifugation (10 min, 3900 × g, 10 °C). Supernatant was decanted and cell pellets stored at -80 °C prior to processing. RNA was extracted using an RNeasy Mini Kit (Qiagen) according to manufacturer’s instructions. Samples were treated with DNaseI (Fermentas) following manufacturer’s instructions but excluding those samples with a 260/280 nm ratio of <2. cDNA was generated using the ImProm-II Reverse Transcriptase System (Promega) or SuperScript IV Reverse Transcriptase System (Invitrogen) with control reactions lacking reverse transcriptase prepared in parallel.

Transcript abundance was determined using primers 1 and 2 for *mntS*, 3 and 4 for *fepD*, 5 and 6 for *rcnA*, 7 and 8 for *nikA*, 9 and 10 for *znuA*, 11 and 12 for *zntA*, 13 and 14 for *copA*, 15 and 16 for *rpoD*, 17 and 18 for *gyrA* each pair designed to amplify ∼100 bp (Supplementary Table 1). qPCR was performed in 20 µl reactions containing 5 ng of cDNA, 400 nM of each primer and PowerUP SYBR Green Master Mix (Thermo Fisher Scientific). Three technical replicates of each biological sample were analysed using a Rotor-Gene Q 2plex (Qiagen, Rotor-Gene-Q Pure Detection Software). Control reactions without cDNA template (qPCR grade water used instead) were run for each primer pair and –RT control reactions were run for the reference gene primer pair (*rpoD* or *gyrA*). qPCR was performed on n = 3 biological replicates for each treatment other than samples treated with 1 mM EDTA for 60 min, with primers specific to *znuA* and *rcnA*, where n = 5 biological replicates were analysed. Where an analysis was run more than once a mean of the determined *C*_q_ values was used subsequently. *C*_q_ values were calculated with LinRegPCR after correcting for amplicon efficiency^14^, with each primer pair and treatment condition considered as an amplicon. Samples where the *C*_q_ value for no template or –RT control was <10 (to the nearest integer) compared to the cDNA containing sample were rejected. These samples were either rerun or DNaseI treatment and cDNA synthesis performed again on RNA samples before running qPCR.

**Table 1.** Calculated available metal concentrations in conditional cells.

| Metal | Sensor | Aerobic<br>[Metal]<sub>buffered</sub> (M) | Anaerobic<br>[Metal]<sub>buffered</sub> (M) | H<sub>2</sub>O<sub>2</sub> treated<br>[Metal]<sub>buffered</sub> (M) |
| --- | --- | --- | --- | --- |
| Mn<sup>2+</sup> | MntR | $6.8(\pm 4) \times 10^{-6}$ | $9.6(\pm 3) \times 10^{-6}$ | $8.8(\pm 2) \times 10^{-5}$ |
| Fe<sup>2+</sup> | Fur | $1.9(\pm 0.2) \times 10^{-6}$ | $6.6(\pm 2) \times 10^{-7}$ | $3.4 \times 10^{-6}$ |
| Co<sup>2+</sup> | RcnR | $6.5(\pm 1) \times 10^{-11}$ | $7.1(\pm 0.5) \times 10^{-11}$ | $2.4 \times 10^{-11}$ |
| Zn<sup>2+</sup> | Zur | $7.0(\pm 1) \times 10^{-12}$ | $6.7(\pm 0.9) \times 10^{-12}$ | $3.2 \times 10^{-11}$ |
| | ZntR | $4.1(\pm 1) \times 10^{-13}$ | $2.9(\pm 0.3) \times 10^{-13}$ | $1.6(\pm 0.4) \times 10^{-13}$ |
| | Mid-point | $1.7 \times 10^{-12}$ | $1.4 \times 10^{-12}$ | $2.2 \times 10^{-12}$ |
| Ni<sup>2+</sup> | NikR | $1.3(\pm 0.7) \times 10^{-13}$ | $8.6(\pm 2) \times 10^{-14}$ | $5.1(\pm 2) \times 10^{-13}$ |
| Cu<sup>+</sup> | CueR | $4.5(\pm 0.6) \times 10^{-20}$ | $1.5(\pm 0.2) \times 10^{-19}$ | $1.1 \times 10^{-20}$ |

### Determination of boundary conditions for the expression of each transcript

Boundary conditions for the calibration of sensor response curves were defined by the minimum and maximum abundance of the regulated transcript. Supplementary Table 2 gives the rationale behind the choice of growth conditions. Cell pellets from cultures exposed to 1.5 and 4 mM FeSO_4_ were brown, difficult to resuspend in RNA extraction buffer, discoloured the buffer, and generated RNA samples with ratios of absorbance at 260/280 nm <2, so were not processed further.

Following completion of all qPCR reactions qPCR data for the control gene (*rpoD*) in each sample was reanalysed in a single LinRegPCR analysis along with collated data for *rpoD* expression in control conditions. The difference between *rpoD C*_q_ in the condition of interest and control condition was determined (average *rpoD C*_q_ for control condition minus average *rpoD C*_q_ for treated sample) (Supplementary Table 3). All aerobic and anaerobic samples were compared against untreated aerobic 2 h controls and anaerobic samples were additionally compared against the most relevant anaerobic control condition (2 or 3 h treatment). Where a difference of >2 was found for *rpoD C*_q_ values, data was either not further analysed or investigated using a second control gene (*gyrA*, Supplementary Table 4), before deciding whether or not to proceed. Notably, cells cultured in the presence of 1 mM ZnSO_4_ showed a significant increase in *rpoD* transcripts relative to the control condition (Supplementary Table 3), which was not observed with *gyrA* (Supplementary Table 4), and these samples were not used further. Likewise, this was the case for samples from cells cultured anaerobically for 3 h either untreated or supplemented with 0.5 mM FeSO_4_ (Supplementary Table 3 and 4). NiSO_4_ treatments where *rpoD C*_q_ values changed by more than two (relative to control condition) and one treatment with EDTA (20 min) were also excluded.

### Intracellular metal availability in conditional cells

The fold change in transcript abundance, relative to the mean of the control condition (lowest expression) for each sensor, was calculated using the 2^-ΔΔCT^ method^15^, with *rpoD* as the reference gene. Fractional responses of metal sensors (*θ*_D_ for de-repressors and co-repressors, *θ*_DM_ for activators) were calculated via equations 1 and 2 calibrating sensor fractional occupancy between 0.01 and 0.99:

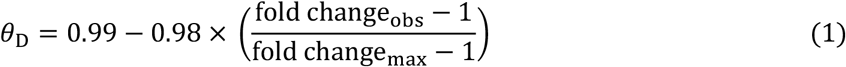

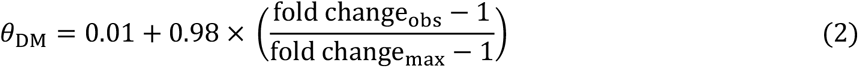

where fold change_obs_ is the fold change in the condition of interest and fold change_max_ is the maximum observed fold change.

Fractional responses of sensors were converted to available metal concentrations using excel spreadsheet (Supplementary Dataset 1) and MATLAB code (Supplementary Note 3) from Osman and coworkers^10^, along with known sensor metal affinities, DNA affinities and protein abundances determined for *Salmonella* sensors (Osman and coworkers^10^), with numbers of DNA binding sites for *E. coli* sensors (Supplementary Table 5). Sensor response curves were also determined with these values and materials.

Intracellular available Δ*G*_Metal_ was calculated using equation 3:

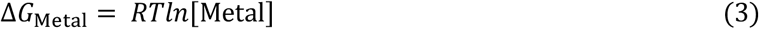

where [Metal] is the intracellular available metal concentration, *R* (gas constant) = 8.314 × 10^−3^ kJ K^-1^ mol^-1^ and *T* (temperature) = 298.15 K.

### Simulated metalation of molecules under bespoke conditions

Three calculators have been created for prediction of molecule metalation in JM109 (DE3) under aerobic and anaerobic conditions, and in response to H_2_O_2_ treatment accounting for multiple inter-metal competitions plus competition from the intracellular buffer as described by Young and coworkers^11^, (https://mib-nibb.webspace.durham.ac.uk/metalation-calculators/ and Supplementary Spreadsheets 1-3). Estimations of molecule metal affinities were taken from cited references.

The web-based calculators were created by converting the Excel spreadsheet created by Young and coworkers^11^ into HTML and JavaScript code that could be run in a web browser. This code was then turned into a plugin for Wordpress, to allow Wordpress page authors to create calculators with different default values for metal availabilities. The source code alone underlying the operation of the web-based calculators has additionally been made available on GitHub and Zenodo^16^.

## Results

### Identifying fold change in abundance of *mntS, fepD, rcnA, nikA, znuA, zntA* and *copA* transcripts by qPCR

We previously defined the range of buffered metal concentrations over which each metal sensor responds on selected promoters (response curves) based on their DNA- and metal-binding affinities, protein molecules per cell under low and high metal exposures, and number of promoter targets within the cell^10^. More recently the position on this response curve under bespoke conditions was used to determine Co^2+^ availability in *E. coli* cells engineered to produce vitamin B_12_^11^. To calculate metal availability under specific conditions, minimum transcript abundance observed by qPCR was first defined to enable fold change in expression under specific conditions to be related to this boundary condition. Calculations of metal availability secondarily require identification of maximum transcript abundance by qPCR to define the opposite boundary.

To select conditions for RNA isolation, JM109 (DE3) was cultured in metals and EDTA to identify ∼15% growth inhibition relative to untreated cells after 2 h exposure (Supplementary Table 6). Where necessary more inhibitory concentrations were used along with shorter or longer exposure times. Anaerobic culture, exposure to H_2_O_2_ and to the Ni-specific chelator DMG followed established protocols^17,18^. Control gene, *rpoD*, was used throughout plus *gyrA* to validate or eliminate samples where *rpoD* expression (*C*_q_) changed by more than two relative to control conditions (Supplementary Tables 3 and 4).

Expression from the *nikA* promoter is repressed in response to rising Ni^2+^ availability by NikR but dependent upon activation by Fnr under anaerobic conditions^19-22^. We observed Ni^2+^ dependent regulation of *nikA* expression both aerobically and anaerobically (Supplementary Table 7). Boundary conditions were therefore independently defined for anaerobically and aerobically grown cells.

Supplementary Tables 7-13 show all of the resulting Δ*C*_q_ values. Change in gene expression relative to the lowest abundance for each transcript was calculated as a series of ΔΔ*C*_q_ values (Supplementary Tables 14-21). In turn ΔΔ*C*_q_ values were expressed as fold increase in gene expression (Fig. 1).

**Figure 1.**
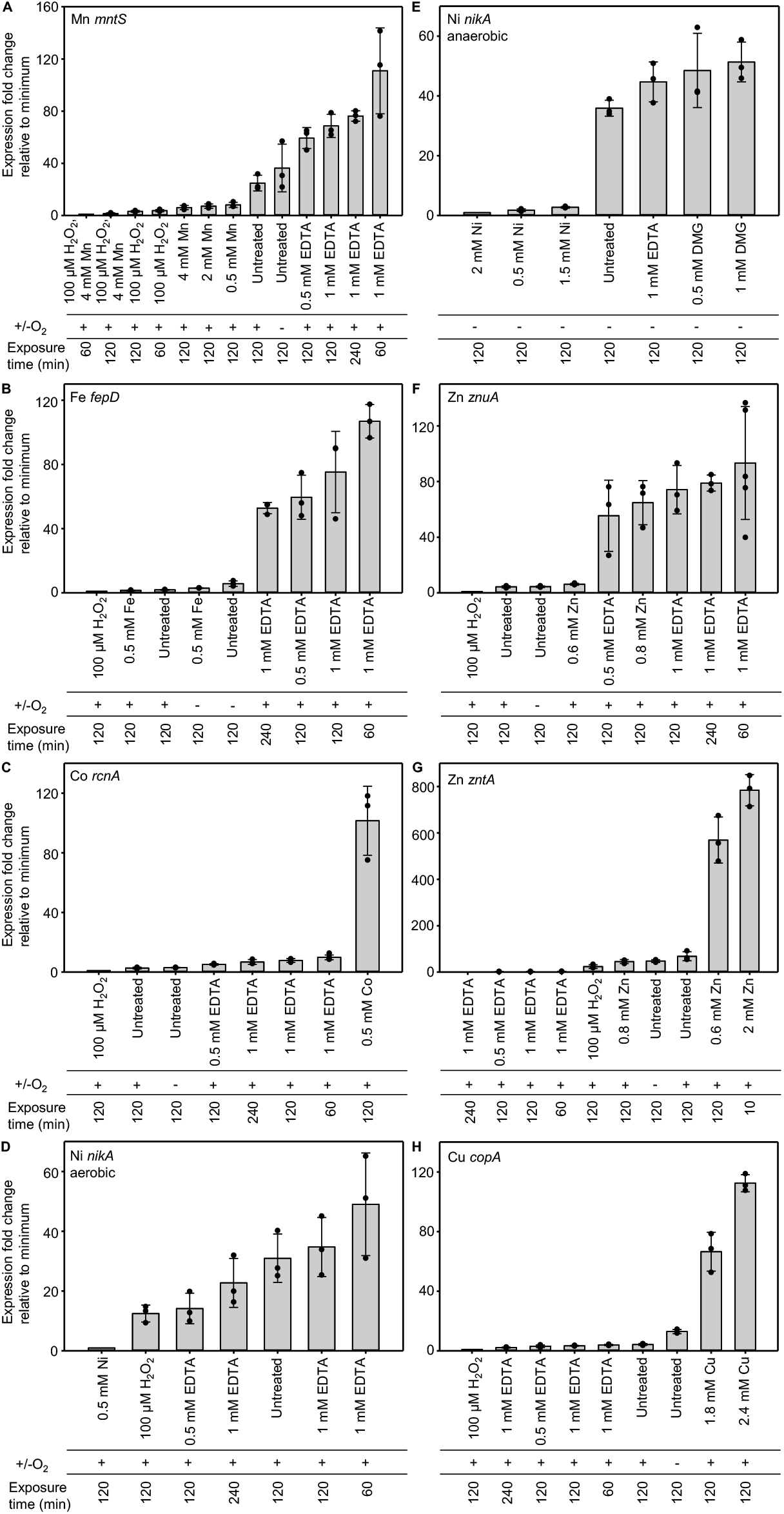
Fold change in metal responsive transcript abundance relative to a minimum. Abundance of seven metal-responsive transcripts relative to the first (left-most) condition shown on each chart (assigned a value of 1) as determined by qPCR. **A**. *mntS* regulated by Mn2+ sensor MntR. **B**. *fepD* regulated by Fe2+ sensor Fur. **C**. *rcnA* regulated by Co2+ sensor RcnR. **D**. *nikA* regulated by Ni2+ sensor NikR (aerobic growth). **E**. *nikA* regulated by Ni2+ sensor NikR (anaerobic growth). **F**. *znuA* regulated by Zn2+ sensor Zur. **G**. *zntA* regulated by Zn2+ sensor ZntR. **H**. *copA* regulated by Cu+ sensor CueR. ΔΔ*C*q values from which these fold changes

### Calibrated responses of MntR, Fur, RcnR, NikR, Zur, ZntR and CueR to Mn^2+^, Fe^2+^, Co^2+^, Ni^2+^, Zn^2+^ and Cu^+^

The fold changes in gene expression shown in Figure 1 need to be calibrated to DNA occupancies (conditional *θ*_D_ for repressors or *θ*_DM_ for activators) and in turn DNA occupancies related to buffered metal concentration. Minimum transcript abundance defines maximum DNA occupancy for co-repressors and de-repressors (*θ*_D_ assigned as 0.99) while defining minimum DNA occupancy for metalated activators (*θ*_DM_ assigned as 0.01). To set the opposite end of the response range for the co-repressors and de-repressors (*θ*_D_ = 0.01) and for activators (*θ*_DM_ =0.99) we selected the highest expression from Figure 1. Conditional *θ*_D_ or *θ*_DM_ values were calculated based on the proportion of the maximum fold change observed for each promoter using common equations for repressors (Equation 1) and a separate relationship for activators (Equation 2).

The relationships between intracellular metal availabilities and *θ*_D_ or *θ*_DM_ for each sensor were calculated essentially as described previously using measured metal affinities, DNA affinities, protein abundance, numbers of DNA targets (Supplementary Table 5), via Excel spreadsheet (Supplementary Dataset 1) available in Osman and coworkers^10^. Notably, the numbers of target promoters for CueR and NikR in *E. coli* differ from *Salmonella* (Supplementary Table 5). *θ*_D_ and *θ*_DM_ values at 0.01 and 0.99 were related to available metal concentration as above and using MATLAB code (Supplementary Note 3) available in Osman and coworkers^10^ (red symbols on Fig. 2).

**Figure 2.**
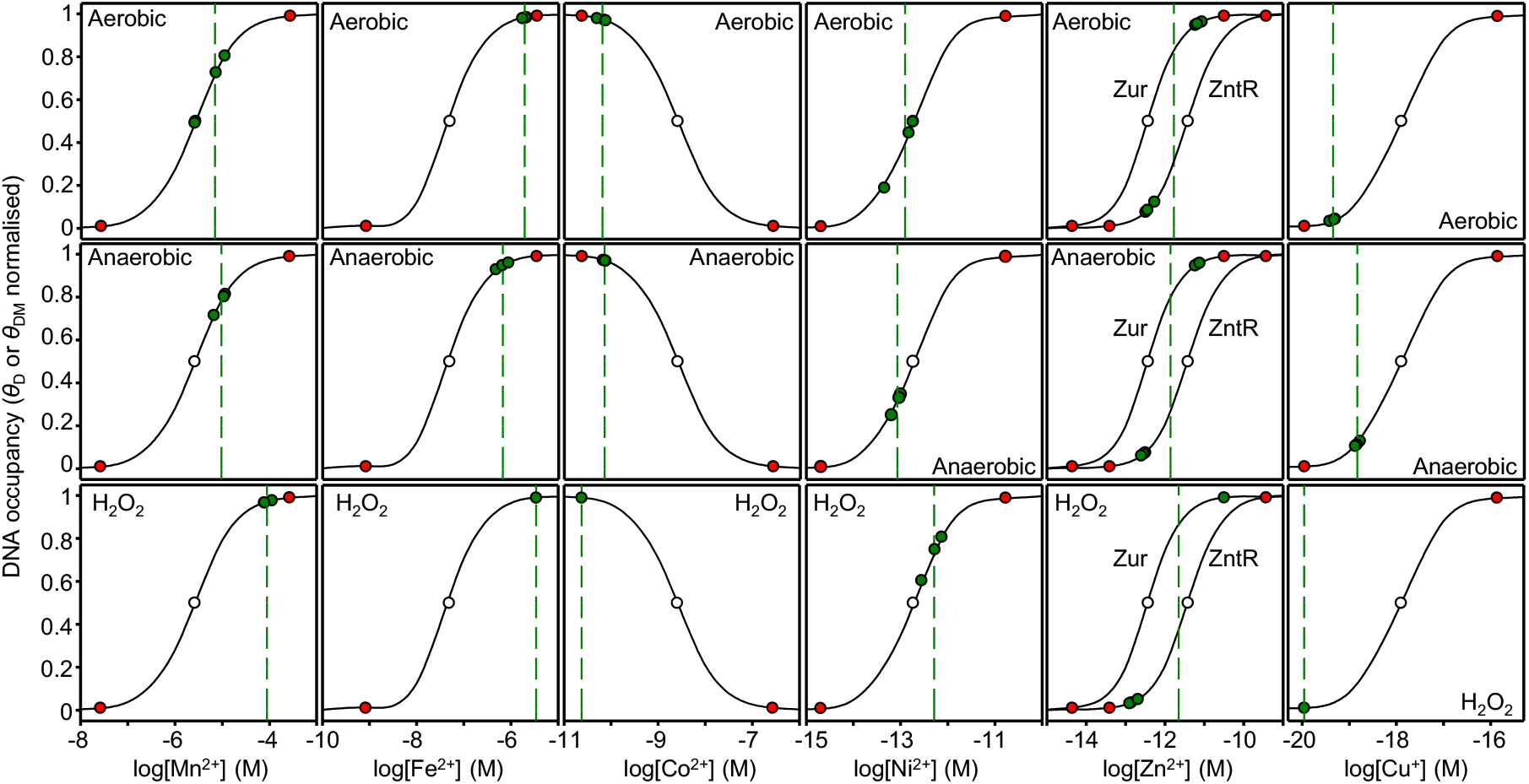
Calibrated responses of metal sensors as a function of intracellular metal-availability. The calculated relationship between intracellular metal availability and normalised DNA-occupancy (*θ*_D_) or for activators normalised DNA-occupancy with metalated sensor (*θ*_DM_). The dynamic range within which each sensor responds to changing intracellular metal availability has been defined as *θ*_D_ or *θ*_DM_ of 0.01 to 0.99 representing the boundary conditions for maximum or minimum fold-changes in transcript abundance (red circles). Mid-point of each range (open circle), replicate values calculated for LB media (green circles), and mean availability (green dashed lines). For Zn^2+^ two curves represent the responses of *znuA* and *zntA*, here the green dashed line is a midpoint between the means for each sensor. Buffered concentrations and curves were calculated using excel spreadsheet (Supplementary Dataset 1) and MATLAB code (Supplementary Note 3) along with thermodynamic properties of sensors described by Osman and coworkers^10^, and numbers of DNA binding sites for *E. coli* sensors (Supplementary Table 5).

### Availabilities of Mn^2+^, Fe^2+^, Co^2+^, Ni^2+^, Zn^2+^ and Cu^+^ in cells cultured in LB media

To estimate the buffered concentrations of available metal inside *E. coli* strain JM109 (DE3) cells grown in LB media, fold changes in abundance of transcripts encoded by the seven metal-responsive genes were calculated in RNA isolated from cells grown aerobically, anaerobically and in response to H_2_O_2_. Values were converted to DNA occupancies for the respective metal-sensor proteins (*θ*_D_ and *θ*_DM_) (green symbols on Fig. 2). Buffered concentrations of available metals were thus derived from their established relationships to *θ*_D_ or *θ*_DM_ for each sensor using MATLAB code (Supplementary Note 3) available in Osman and coworkers^10^ (Table 1).

### Free energies for forming half-saturated metal-complexes at intracellular metal availabilities in *E. coli*

Inside cells available metals are mostly bound to labile sites rather than fully hydrated. For clarity, and to assist subsequent data manipulations, metal availabilities were expressed in terms of free energies for forming complexes with proteins (or other types of molecules) that will be 50% metalated at the respective buffered available metal concentrations shown in Table 1. The dissociation constant (*K*_D_) of such a molecule matches the metal concentration and free energy (*ΔG*) is calculated using the relationship shown in Equation 3. These data reveal how tightly labile metals are bound and hence the magnitude of competition for each metal inside *E. coli* JM109 (DE3) under specific growth conditions (red symbols on Fig. 3). Standard deviations were calculated based upon the triplicated determinations of buffered concentration which have been averaged in Table 1. For comparison, previously used metal-availabilities at the mid-points of the ranges of each sensor for each metal in idealised cells, are also shown (grey symbols in Fig. 3).

**Figure 3.**
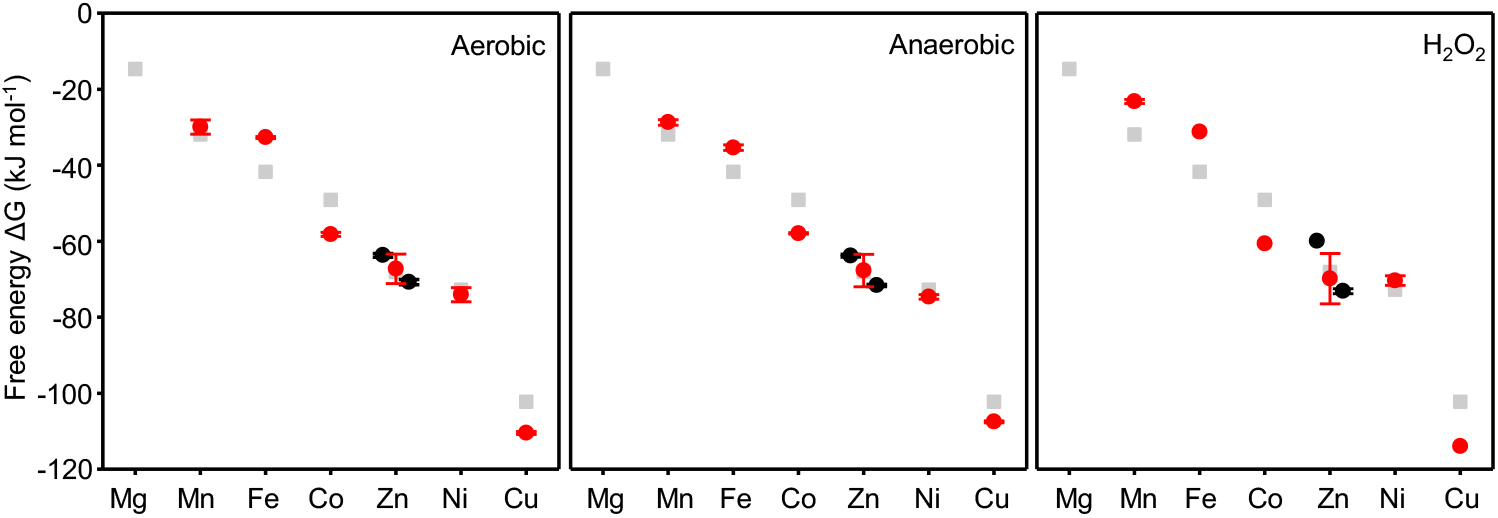
Free energies of available metal in *E. coli* JM109 (DE3). Intracellular available free energies for metal binding to molecules that would be 50 % saturated at the available metal concentration (Table 1) in conditional cells (red circles with standard deviation). For Zn^2+^ separate values have been derived based on the response of Zur and ZntR (black circles with standard deviation, Zur = left, ZntR = right). Grey squares are values for idealised cells reflecting the mid-point of sensor ranges as used in previous versions of calculators.

### Three web-based metalation calculators

To predict the metalation states of proteins of known metal affinities, three metalation calculators were developed based on the intracellular metal availabilities estimated in cells grown in LB media under aerobic, anaerobic, and H_2_O_2_ exposed, conditions (Fig. 3). Initially calculators were produced as spreadsheets (Supplementary Spreadsheets 1-3), as described previously^11^. The spreadsheets complete two operations, firstly calculating the difference in free energy for metal binding to the protein versus competing sites of the intracellular milieu, secondly accounting for inter-metal competition as described previously^11^. Web-based versions of the three calculators have been generated, each pre-populated with the metal availabilities determined under each of the three growth conditions (https://mib-nibb.webspace.durham.ac.uk/metalation-calculators/). Toggle switches exclude metals from the calculations enabling simulations for proteins where some affinities are unknown (toggle switches are visible on the far left in Figure 4). Metal affinities are entered as dissociation constants, *K*_D_, and the calculators output free energies for forming the respective metal complexes as well as occupancies.

**Figure 4.**
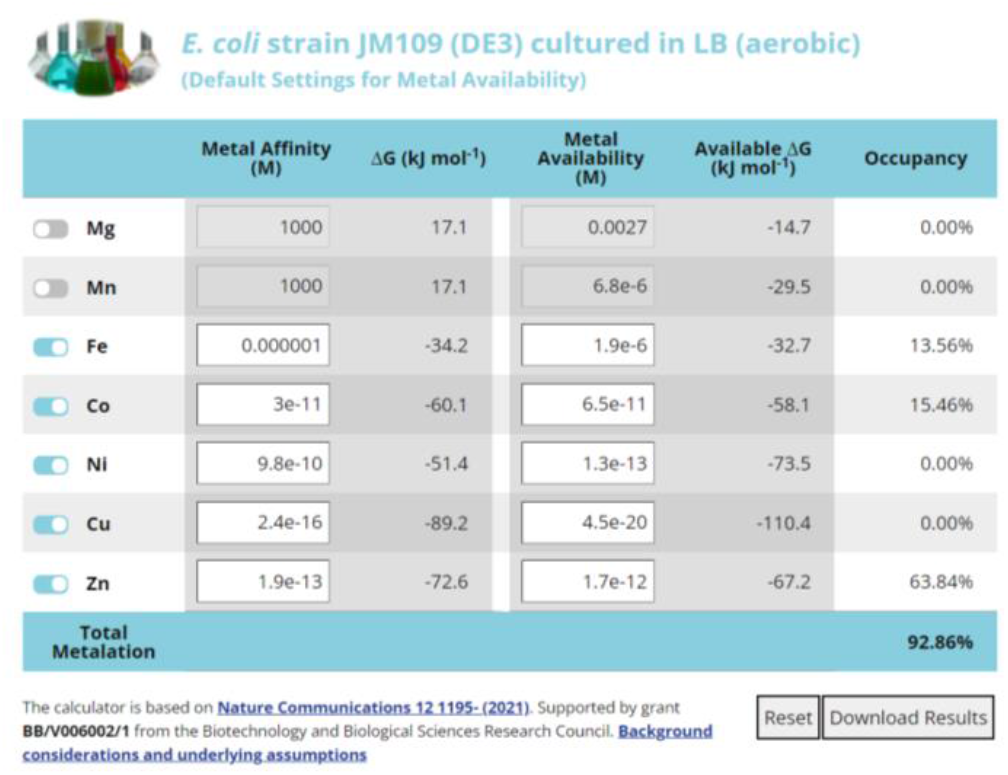
Web based metalation calculators. Screenshot of one of the three metalation calculations available online (https://mib-nibb.webspace.durham.ac.uk/metalation-calculators/). These calculate metal occupancy of a protein or other molecule whose affinities have been entered into the first column of editable cells. Toggle switches exclude metals for which affinities are unknown (left) and effects of other metal availabilities can be simulated by entering concentrations into the second column of editable cells.

## Discussion

### Metal availability follows the Irving-Williams series under all three conditions

Metals tend to associate with nascent proteins with the following order of preference (from weakest to tightest): Mg^2+^ < Mn^2+^ < Fe^2+^ < Co^2+^ < Ni^2+^ < Cu^2+^ (Cu^+^) > Zn^2+^, as set out in the original Irving Williams series and noting the subsequent addition of monovalent copper^3,6^. The availabilities of these metals (as divalent forms except copper which is predominantly monovalent in the cytosol) in *E. coli* JM109 (DE3) follows this series under all three conditions (Fig. 3, Table 1). Notably the series is ambiguous about the exact position of Zn^2+^, only specifying that binding is weaker than copper^3^, and in conditional cells the free energy for forming complexes with available Zn^2+^ is indeed less negative than copper, but also slightly less negative than Ni^2+^ (Fig. 3, Table 1). By maintaining the available forms of the tightest binding metals at the lowest free energies for forming metal complexes the challenge to correctly metalate proteins in cells is substantively overcome^3,7,10,11^.

### In LB media Fe^2+^ is more, and Cu^+^ and Co^2+^ less, available than in idealised cells

Comparisons of the calculated free energies for forming complexes with available metals in the cytosol of idealised cells (grey symbols on Fig. 3) with those inside conditional *E. coli* JM109 (DE3) (red symbols on Fig. 3) reveals that Fe^2+^ is more available than previously suggested^10^. Notably, the availability of Fe^2+^ is only slightly less than Mn^2+^ (Fig. 3, Table 1), and Fur dependent expression approaches, or defines, the upper boundary condition (Fig. 2). In some bacteria Fe^2+^ sensors have been discovered that detect highly elevated Fe^2+^, including most recently riboswitches^23,24^. These observations raise the intriguing possibility that Fe^2+^ availability may become even greater than detected here, perhaps even exceeding the availability of Mn^2+^. Enhanced binding of Fe^2+^ Fur has been documented at some *E. coli* promoters under anaerobic conditions, but not the *fepD* promoter^25^. Moreover, the number of Fur molecules per cell is insufficient to populate all Fur-target promoters in *Salmonella* (and by inference *E. coli*) cells grown aerobically but becomes sufficient after iron supplementation, and hence some target promoters with weaker affinities for Fur could remain vacant in *E. coli* cells cultured aerobically in LB media^10^. Calibration of a different Fur-target promoter could allow the detection of some further increase in available Fe^2+^ in anaerobically cultured cells, and iterative improvements in estimated metal availabilities will be updated on the web-based versions of the calculators.

The number of promoter targets for CueR and NikR differ in *E. coli* relative to *Salmonella* (Supplementary Table 5). The modelled responses of these sensors are consequently altered in *E. coli* slightly changing availabilities in idealised cells as well as conditional cells (Fig. 2 and 3). Previous data established in another strain of *E. coli* that the availability of Co^2+^ was considerably less than predicted in idealised cells and this observation is confirmed here for JM109 (DE3)^11^. The availability of copper is also significantly less than in idealised cells, defining the lower boundary condition in response to H_2_O_2_ (Fig. 2 and 3). Copper is an extremely potent pro-oxidant catalysing the Fenton reaction and a reduction in available Cu^+^ in the presence of H_2_O_2_ will limit production of the deadly hydroxyl radical. The free energy of exchangeable and available Ni^2+^ does not significantly change under anaerobic conditions where it is known that the metal is imported to supply hydrogenase^26^. In anaerobic cells the magnitude of additional import by the Nik-system and the magnitude of flux through the Hyp-metallochaperones into nascent protein, notably hydrogenase, may be approximately matched.

### H_2_O_2_ increases Mn^2+^ availabilities

Mn^2+^ is an effective antioxidant^27^. Detection of H_2_O_2_ by OxyR triggers Mn^2+^ import and activates manganese superoxide dismutase (SodA) with evidence that SodA is otherwise mis-metalated with iron and inactive^28,29^. Here we similarly detect elevated Mn^2+^ in response to H_2_O_2_ (Fig. 3 and Table 1). Some treatments, including potentially exposure to H_2_O_2_, may directly modify the sensors independently of effects on metal availability, raising a caveat that readouts of free energies for metalation may become less accurate for some metals under these conditions.

### Simulated metalation of *E. coli* HypB, SodA and selected riboswitches

The HypB metallochaperone recruits Ni^2+^ to the HypA/B complex and subsequently to hydrogenase in *E. coli* grown under anaerobic conditions^30^. Based on a Ni^2+^ affinity of 6 × 10^− 14^ M and Zn^2+^ affinity of 2.2 × 10^−11^ M reported by Zamble and co-workers^31^, and excluding metals other than Ni^2+^ or Zn^2+^ (or using arbitrary weak HypB affinities for other metals in the spreadsheet), the anaerobic calculator (https://mib-nibb.webspace.durham.ac.uk/metalation-calculators/, Supplementary Spreadsheet 2) reassuringly predicts HypB to be predominantly metalated by Ni^2+^ in this condition (57.4% Ni^2+^, 2.6% Zn^2+^).

Metals are kinetically trapped within SodA but attempts have been made to measure affinities via thermal unfolding of metalated and un-metalated protein^32^. This approach generates affinities of 3.1 × 10^−9^ M for Mn^2+^ and 2.5 × 10^−8^ M for Fe^2+^ ref.^32^. SodA becomes correctly metalated after exposure to H_2_O_2_ and indeed the H_2_O_2_ calculator predicts 99.5% occupancy with Mn^2+^ and 0.5% occupancy with Fe^2+^ in this condition. However, using these affinities the aerobic calculator predicts only slightly reduced occupancy to 96.6% with Mn^2+^ and only slightly increased to 3.4% occupancy with Fe^2+^. The reported affinities do not follow the Irving-Williams series raising a tantalising possibility that they do not reflect flexible sites at which exchangeable metals partition onto the nascent unfolded SodA protein prior to kinetic trapping. Simulations using an affinity for Mn^2+^ which is 10-fold weaker than the reported affinity for Fe^2+^, abiding by the Irving-Williams series, do flip occupancy from predominantly Fe^2+^ in the absence of H_2_O_2_ to predominantly Mn^2+^ in the presence of H_2_O_2_, by using the respective two calculators (26.1% Mn^2+^ and 72.9% Fe^2+^ aerobically to 72.0% Mn^2+^ and 27.8% Fe^2+^ in H_2_O_2_).

A spectroscopically active variant of a *czcD* riboswitch, previously designated as a Co^2+^/Ni^2+^ sensor, was recently shown to respond to Fe^2+^ when studied in *E. coli*^23^. This experimental observation was shown to align with predictions from calculations based on idealised cells and the thermodynamically calibrated ranges of *Salmonella*, and by inference *E. coli*, metal sensors^10,23^. The predictions are also supported here when using the aerobic calculator to give occupancies of 74.6% Fe^2+^, negligible occupancy with either Ni^2+^ or Co^2+^ (0.01% Co), also zero occupancy with Zn^2+^ and a modest 9.7% occupancy with Mn^2+^. An alteration in the documented selectivity of this riboswitch from Ni^2+^ and Co^2+^ to Fe^2+^ seems apposite.

### Uses, constraints and future prospects for metalation calculators

Predictions of the calculators can be used to: identify and/or confirm metal-specificity; infer mis-metalation; reveal erroneous metal affinity measurements; suggest where metal availabilities may differ from calculator values; indicate where additional mechanisms assist metalation. Several of these are exemplified by simulations in the preceding section.

The calculators assume that the molecule of interest does not deplete the buffered available metal pool. This suggests a potential constraint associated with the widespread use of *E. coli* overexpression systems to generate recombinant proteins for research purposes and here the outputs of metal sensors might reveal where overexpression of a metalloprotein depletes available metals. The calculators also assume a 1:1 metal complex with a preformed metal site on the molecules of interest. They cannot directly predict occupancies for metal-dependent assemblies in which the ligands are derived from more than one molecule: Notably, the degree of metal binding to such complexes will depend on the intracellular ligand concentrations and hence prone to occur when proteins are overexpressed in *E. coli*. It is anticipated that future iterations of the calculator could be developed for such metal-dependent assemblies where affinities are reported as *β* values in M^-2^.

As noted earlier some strains of *E. coli* commonly used for protein overexpression are aberrant in metal homeostasis. For example, strain BL21 (DE3) lacks the *rcn* genes, is mutated in *fnr* and lacks the *mod* operon for molybdate uptake^33^. It is anticipated that bespoke versions of the calculator might be generated for such strains. Additionally, *E. coli* unlike *Salmonella* lacks a dedicated Co^2+^ uptake system^34^. Under- or mis-metalation of Co^2+^ or Ni^2+^ (and indeed molybdenum) requiring proteins has been documented and may be common following overexpression in *E. coli*^*11,35*^.

The calculators predict metal occupancies based upon a population of intracellular metal-buffering sites at thermodynamic equilibrium with the molecule of interest. The magnitude of disparities from these predictions can thus be used to establish the magnitude of additional contributions to metal-specificity, for example where the distribution of metalated products is, at least in part, kinetically determined. Localisation of a nascent metalloprotein proximal to a metal importer may favour metalation in a niche where metal availability is greater than average for the compartment: There is also limited evidence of direct metal transfer by ligand exchange from importers to docked-metalloproteins^36-39^. Kinetic bias could theoretically occur where a buffering molecule preferentially accesses a specific metal site with dedicated metallochaperones representing an extreme example.

Previous use of an earlier iteration of a metalation calculator revealed that the Co^2+^-chaperone CobW alone would be unable to acquire Co^2+^ in *E. coli*^11^. Binding of GTP and Mg^2+^ increased the Co^2+^ affinity sufficiently to enable metalation. In contrast, after hydrolysis, it was previously noted that binding of GDP would lead to Co^2+^ release illustrating how use of such a calculator can uncover the contributions of crucial molecular interactions to metalation, and hence mechanisms of action^11^.

Simulations for proteins of other organisms but using these calculators for *E. coli* strain JM109 (DE3) (https://mib-nibb.webspace.durham.ac.uk/metalation-calculators/ and Supplementary Spreadsheets 1-3) may indicate where availabilities could depart from those estimated here. It is anticipated that future iterations of these calculators may be generated with metal availabilities established for other cell types, either via bespoke calibration of metal sensors from other strains or through use of other approaches to estimate metal availability. This could include the use of small molecule metal probes or genetically encoded probes of metal availability. Furthermore, subtle differences between the metal sensors of *E. coli* and *Salmonella* might have led to disparities in estimating metal availabilities that could emerge in simulations of metalation of *E. coli* proteins (Supplementary Table 22). This could be resolved by thermodynamically characterising the respective *E. coli* sensor in the manner performed for *Salmonella*^*10*^, the web-based version of the calculators will catalogue such updates in estimates of metal availability.

There is a view that many reported metal affinities of proteins are not correct, and a recent article provides a guide to such measurements^40^. Simulations using the metalation calculators might predict aberrant metalation that is indicative of erroneous values. Separately, the kinetic trapping of metals during folding can abide by the Irving-Williams series, as documented for the Mn^2+^ cupin MncA^41^. As noted earlier, the challenge to measure the relevant affinities along the folding pathway when a metal becomes kinetically trapped is especially interesting.

Metal binding to some proteins, such as the Mn^2+^ form of ribonucleotide reductase, subtlety depart from the Irving-Williams series, binding Mn^2+^ in preference to Fe^2+^ ref.^42^. This is facilitated by redox changes and the introduction of steric selection via cooperativity at the di-metal site^42^. An inference might be that Mn^2+^ ribonucleotide reductase is metalated under conditions where Mn^2+^ is not more available than Fe^2+^: notably this state is approached in Figure 3 and perhaps might be detected in anaerobically grown cells after calibration of a weaker Fur-regulated promoter. Synthetic proteins have been designed which do not select metals according to the order of metal-selectivity given by the Irving-Williams series and as commonly observed in nature^43^. With some analogy to ribonucleotide reductase, this is achieved through the introduction of steric selection through the cooperative binding of multiple metals. This raises questions as to whether evolution could have achieved greater selectivity bypassing the necessity to maintain metal availabilities as the inverse of the Irving-Williams series. It has been suggested that such enhanced selectivity comes at the price of increased rigidity and an impaired catalytic landscape^43^. Importantly, for artificial metalloproteins to acquire the correct metal in a cell, engineered selectivity need only be sufficient to confer correct metalation as predicted by metalation calculators based on the prevailing intracellular thermodynamic landscape.

In industrial biotechnology there is potential for a mismatch between introduced heterologous proteins and metal availabilities in organisms used for bioprocessing and biotransformation. This is illustrated by the calculated mis-metalation of *Rhodobacter* CobW with Zn^2+^ in *E. coli* engineered to make vitamin B_12^11^_. Outputs of a metalation calculator thus present solutions for optimising sustainable manufacturing, in this example by chelating Zn^2+^ from fermentation media, supplementing fermentation media with Co^2+^, or by engineering the respective homeostatic systems for Co^2+^ and/or Zn^2+^ in *E. coli*. With so many enzymes requiring metals this offers considerable potential for exploitation in the transition to more sustainable bio-based **manufacturing**.

## Acknowledgements

This work was supported by Biotechnology and Biological Sciences Research Council awards: BB/V006002/1, BB/S009787/1, BB/R002118/1, BB/S014020/1, a Royal Commission for Exhibition of 1851 Research Fellowship (TRY). We thank Deenah Osman, Martin Warren, Peter Chivers, William Michaels and Maria Martini for discussions and supporting intellectual contributions.

