## Supplementary Data for "Metalation calculators for *E. coli* strain JM109 (DE3): Aerobic, anaerobic and hydrogen peroxide exposed cells cultured in LB media"

| No. | Name | Sequence | Product size (bp) | Reference |
|---|---|---|---|---|
| 1 | *mntS*_F | 5’-GTATGCGCGTGTTTAGTCATTC-3’ | 105 | This work |
| 2 | *mntS*_R | 5’-TATCGGAAGGTTTATCTTGCTG-3’ | 105 | This work |
| 3 | *fepD*_F | 5’-TGCAAACCCTCACCCGAAAC-3’ | 111 | This work |
| 4 | *fepD*_R | 5’-GCGCGGAAGAGTAACCAAACAG-3’ | 111 | This work |
| 5 | *rcnA*_F | 5’-GAACCAGGGCACTCAAAAAC-3’ | 108 | ref.[1] |
| 6 | *rcnA*_R | 5’-TGCGGTATGCGAAATAGTTG-3’ | 108 | ref.[1] |
| 7 | *nikA*_F | 5’-AACCCGCACCTTTACACGCC-3’ | 114 | This work |
| 8 | *nikA*_R | 5’-AGTCCAGCTTTTTGCCAGCC-3’ | 114 | This work |
| 9 | *znuA*_F | 5’-GTTTGGACTGACACCGCTTG-3’ | 111 | ref.[2] |
| 10 | *znuA*_R | 5’-ACGCAGGTTGCTTTTTGCTC-3’ | 111 | ref.[2] |
| 11 | *zntA*_F | 5’-CGAAGCACAGGTTGCTGAAC-3’ | 114 | This work |
| 12 | *zntA*_R | 5’-CCGGCAGCGCAAATCAATAC-3’ | 114 | This work |
| 13 | *copA*_F | 5’-GTCACAAACTATCGACCTGACCC-3’ | 104 | This work |
| 14 | *copA*_R | 5’-CATCCGCCTGCTCAACATCC-3’ | 104 | This work |
| 15 | *rpoD*_F | 5’-GTGGCTTGCAGTTCCTTGAC-3’ | 108 | ref.[2] |
| 16 | *rpoD*_R | 5’-AGGTTGCGTAGGTGGAGAAC-3’ | 108 | ref.[2] |
| 17 | *gyrA*_F | 5’-AGGGCTGATGGAACACATCC-3’ | 108 | This work |
| 18 | *gyrA*_R | 5’-ATATACACCTTGCCGCGACC-3’ | 108 | This work |

**Supplementary Table 1**. qPCR primers used in this work.

| +/- $O_2$ | Treatment | Exposure time (min) | Concentration (mM) | Rationale |
|---|---|---|---|---|
| + | Mn | 120 | 0.5, 2, 4 | No growth inhibition observed upon Mn treatment, indeed higher concentrations of Mn appeared to enhance growth. A range of Mn concentrations from the growth experiment were sampled. |
| + | $H_2O_2$ | 60, 120 | 0.1 | *mntH* (Mn import) expression is upregulated by OxyR in response to $H_2O_2$ treatment. Maximal induction observed following 45-60 min treatment with 100 μM $H_2O_2$ in *Salmonella* and similar results observed in *E. coli* [3,4]. Hypothesised that this treatment could increase intracellular Mn availability more than exposure to Mn alone. |
| + | $H_2O_2$ / Mn | 60, 120 | 0.1 / 4 | Hypothesised that $H_2O_2$ exposure in the presence of Mn might give maximum intracellular Mn and therefore lowest expression of *mntS*. |
| + | EDTA | 120 | 0.5 | 16% growth inhibition relative to untreated cultures. |
| + | EDTA | 20, 60, 120, 240 | 1 | To minimise metal availability in growth medium increased [EDTA] to 1 mM and varied exposure times. 1 mM EDTA previously used for calibration of Co and Zn availability in *E. coli* producing $B_{12}$[2]. |
| + | Fe | 120 | 0.5 | 19% growth inhibition relative to untreated cells. |
| + | Fe | 120 | 1.5 | Increased [Fe] to maximise intracellular Fe availability. |
| + | Fe | 120 | 4 | Increased [Fe] to maximise intracellular Fe availability. |
| + | Co | 120 | 0.5 | 17% growth inhibition relative to untreated cells. Obtained a similar maximum fold induction of *rcnA* to that observed previously[2]. |
| + | Ni | 120 | 0.5 | Here a growth condition was selected which gave negligible growth inhibition relative to untreated cells (3%) due to an initial misinterpretation of growth data. However, comparison of *rpoD Cq* for Ni exposed cultures with the control condition suggest that this is the maximum permissible [Ni] for this analysis (Supplementary Table 3). |
| + | Ni | 120 | 2 | Increased [Ni] to maximise intracellular Ni availability. |
| + | Ni | 120 | 3 | Increased [Ni] to maximise intracellular Ni availability. |
| + | Zn | 120 | 0.6 | 21% growth inhibition relative to untreated cultures. |
| + | Zn | 60, 120 | 1 | Increased [Zn] to maximise intracellular Zn availability. |
| + | Zn | 120 | 0.8 | ~50% growth inhibition relative to untreated cultures. Included as 60 and 120 min treatment with 1 mM Zn resulted in aberrant expression of *rpoD*. |
| + | Zn | 10 | 1, 2 | Hypothesised that maximum response of *zntA* might be observed in response to 'zinc shock'. |
| + | Cu | 120 | 1.8 | 14% growth inhibition relative to untreated cultures. |
| + | Cu | 120 | 2.4 | Increased [Cu] to maximise intracellular Cu availability. 36% growth inhibition relative to untreated cultures. Aberrant expression of *rpoD* had been observed in response to [Ni] and [Zn] which resulted in >50% growth inhibition relative to untreated cultures so a less inhibitory concentration was chosen here. |
| - | Fe | 120, 180 | 0.5 | Fe availability may be higher under anaerobic conditions[5,6]. Anaerobic sachets used reduce $O_2$ to <1 % within 30 min and <0.1% within 3 h. [Fe] selected based on earlier results. |
| - | Ni | 120 | 0.5 | [Ni] chosen by analogy to aerobic experiments. |
| - | Ni | 120 | 1.5, 2 | Increased [Ni] to maximise intracellular Ni availability with <50% growth inhibition. |
| - | EDTA | 120 | 1 | [EDTA] chosen by analogy to aerobic experiments. |
| - | DMG | 120 | 0.5, 1 | DMG used as Ni specific chelator for maximum expression of *nikA* under anaerobic conditions. 0.5 and 1 mM DMG treatment results in a significant reduction in hydrogenase activity in anaerobic *Salmonella*[7]. |

**Supplementary Table 2**. Rationale for choice of growth conditions for determining sensor calibration boundary conditions.

| +/- $O_2$ | Treatment | Exposure time (min) | *rpoD* $C_q$ difference relative to untreated aerobic samples | *rpoD* $C_q$ difference relative to corresponding (120 or 180 min) anaerobic untreated samples |
|---|---|---|---|---|
| $+O_2$ | 0.5 mM EDTA | 120 | 0.33(±0.5) | n.a. |
| $+O_2$ | 1 mM EDTA | 20 | 2.0(±0.1) | n.a. |
| $+O_2$ | 1 mM EDTA | 60 | -0.56(±0.2) | n.a. |
| $+O_2$ | 1 mM EDTA | 120 | 0.040(±0.2) | n.a. |
| $+O_2$ | 1 mM EDTA | 240 | 0.15(±0.1) | n.a. |
| $+O_2$ | 0.5 mM Mn | 120 | 1.3(±0.3) | n.a. |
| $+O_2$ | 2 mM Mn | 120 | 1.7(±0.3) | n.a. |
| $+O_2$ | 4 mM Mn | 120 | 1.2(±0.09) | n.a. |
| $+O_2$ | 100 µM $H_2O_2$ | 60 | 0.86(0.3) | n.a. |
| $+O_2$ | 100 µM $H_2O_2$ | 120 | 1.5(±0.3) | n.a. |
| $+O_2$ | 100 µM $H_2O_2$ 4 mM Mn | 60 | 1.2(±0.6) | n.a. |
| $+O_2$ | 100 µM $H_2O_2$ 4 mM Mn | 120 | 0.35(±1) | n.a. |
| $+O_2$ | 0.5 mM Fe | 120 | 0.82(±0.1) | n.a. |
| $+O_2$ | 0.5 mM Co | 120 | -0.57(±0.1) | n.a. |
| $+O_2$ | 0.5 mM Ni | 120 | 1.9(±0.6) | n.a. |
| $+O_2$ | 2 mM Ni | 120 | 2.0(±0.1) | n.a. |
| $+O_2$ | 3 mM Ni | 120 | 3.2(±0.2) | n.a. |
| $+O_2$ | 0.6 mM Zn | 120 | 0.50(±0.08) | n.a. |
| $+O_2$ | 0.8 mM Zn | 120 | 0.89(±0.4) | n.a. |
| $+O_2$ | 1 mM Zn | 10 | 2.8(±0.2) | n.a. |
| $+O_2$ | 1 mM Zn | 60 | 2.7(±0.1) | n.a. |
| $+O_2$ | 1 mM Zn | 120 | 3.2(±0.4) | n.a. |
| $+O_2$ | 2 mM Zn | 10 | 1.3(±0.1) | n.a. |
| $+O_2$ | 1.8 mM Cu | 120 | 1.3(±0.1) | n.a. |
| $+O_2$ | 2.4 mM Cu | 120 | 0.97(±0.2) | n.a. |
| $-O_2$ | 0.5 mM Fe | 120 | -0.37(±0.2) | 0.43(±0.2) |
| $-O_2$ | 0.5 mM Fe | 180 | 2.8(±0.3) | 0.17(±0.3) |
| $-O_2$ | 0.5 mM DMG | 120 | 1.3(±0.04) | 1.8(±0.05) |
| $-O_2$ | 1 mM DMG | 120 | 1.4(±0.1) | 1.8(±0.1) |
| $-O_2$ | 1 mM EDTA | 120 | -0.98(±0.4) | -0.50(±0.4) |
| $-O_2$ | Untreated | 120 | -0.82(±0.06) | n.a. |
| $-O_2$ | Untreated | 180 | 2.7(±0.2) | n.a. |
| $-O_2$ | 0.5 mM Ni | 120 | 0.53(±0.8) | 0.98(±0.8) |
| $-O_2$ | 1.5 mM Ni | 120 | -0.50(±0.1) | -0.026(±0.1) |
| $-O_2$ | 2 mM Ni | 120 | 0.49(±0.2) | 0.96(±0.2) |

**Supplementary Table 3**. *rpoD* $C_q$ difference between values determined for treated samples (and anaerobic untreated samples) and the mean value determined for untreated aerobic cultures and, where appropriate, relative to the mean value determined for the corresponding untreated anaerobic cultures. Values shown as mean ± standard deviation.

| +/- $O_2$ | Treatment | Exposure time (min) | *gyrA* $C_q$ difference relative to untreated aerobic samples | *gyrA* $C_q$ difference relative to corresponding (180 min) anaerobic untreated samples |
|---|---|---|---|---|
| +$O_2$ | 1 mM Zn | 120 | 1.5(±0.7) | n.a. |
| -$O_2$ | Untreated | 180 | 0.90(±0.04) | n.a. |
| -$O_2$ | 0.5 mM Fe | 180 | 1.1(±0.2) | 0.24(±0.2) |

**Supplementary Table 4**. *gyrA* $C_q$ difference between values determined for treated samples (and anaerobic untreated samples) and the mean value determined for untreated aerobic cultures plus, where appropriate, relative to the mean value determined for the corresponding untreated anaerobic cultures. Values shown as mean ± standard deviation.

| Sensor | Targets | Total |
| --- | --- | --- |
| MntR | *mntS/R*<br>*mntH*<br>*mntP*<br>*dps* | 4 |
| Fur | *fepD* | 37 |
| RcnR | *rcnA* | 1 |
| NikR | *nikA* | 1 |
| CueR | *copA*<br>*cueO* | 2 |
| ZntR | *zntA* | 1 |
| Zur | *znuABC*<br>*zinT*<br>*L31P*<br>*pliG* | 4 |

**Supplementary Table 5**. List of target genes for *E. coli* metal sensors. Experimentally validated targets for each metal sensor in *E. coli*[8-14]. For Fur the size of the regulon was estimated from the number of transcriptional units with regulatory sequences bound by Fur during exposure to iron-replete, but not iron-starvation conditions[15,16]. *fepD* is shown as a representative target.

| | | | % growth relative to untreated | | | | |
|---|---|---|---|---|---|---|---|
| +/- $O_2$ | Treatment | Concentration (mM) | 20 min | 60 min | 120 min | 180 min | 240 min |
| $+O_2$ | $MnCl_2$ | 0.5 | - | 97(±2) | 105(±11) | 101(±2) | 101(±6) |
| $+O_2$ | $MnCl_2$ | 1 | - | 151(±1) | 155(±5) | 154(±4) | 136(±5) |
| $+O_2$ | $MnCl_2$ | 1.5 | - | 149(±5) | 144(±5) | 145(±7) | 127(±3) |
| $+O_2$ | $MnCl_2$ | 2 | - | 111(±12) | 101(±7) | 108(±7) | 103(±3) |
| $+O_2$ | $MnCl_2$ | 3 | - | 238(±3) | 214(±13) | 191(±7) | 161(±12) |
| $+O_2$ | $MnCl_2$ | 4 | - | 232(±6) | 184(±2) | 171(±11) | 142(±2) |
| $+O_2$ | $FeSO_4$ | 0.5 | - | 91(±1) | 81(±1) | 83(±0) | 89(±0) |
| $+O_2$ | $FeSO_4$ | 1 | - | 69(±2) | 64(±0) | 64(±1) | 67(±4) |
| $+O_2$ | $FeSO_4$ | 1.5 | - | 64(±2) | 59(±3) | 57(±0) | 60(±3) |
| $+O_2$ | $FeSO_4$ | 2 | - | 95(±2) | 98(±3) | 94(±4) | 92(±5) |
| $+O_2$ | $FeSO_4$ | 3 | - | 107(±2) | 84(±9) | 67(±6) | 56(±5) |
| $+O_2$ | $FeSO_4$ | 4 | - | 107(±4) | 85(±2) | 69(±3) | 56(±2) |
| $+O_2$ | $CoCl_2$ | 0.1 | - | 88(±2) | 101(±1) | 97(±2) | 91(±4) |
| $+O_2$ | $CoCl_2$ | 0.5 | - | 78(±0) | 83(±1) | 82* | 74(±1) |
| $+O_2$ | $CoCl_2$ | 1 | - | 57(±2) | 46(±1) | 36(±2) | 32(±1) |
| $+O_2$ | $NiSO_4$ | 0.5 | - | 93(±0) | 97(±5) | 92(±1) | 105(±3) |
| $+O_2$ | $NiSO_4$ | 1 | - | 74(±3) | 95(±7) | 75(±9) | 82(±9) |
| $+O_2$ | $NiSO_4$ | 1.5 | - | 76(±7) | 82(±7) | 68(±2) | 73(±2) |
| $+O_2$ | $NiSO_4$ | 2 | - | 63(±4) | 41(±5) | 40(±9) | 38(±8) |
| $+O_2$ | $NiSO_4$ | 2.5 | - | 60(±2) | 34(±8) | 30(±5) | 21(±2) |
| $+O_2$ | $NiSO_4$ | 3 | - | 50(±15) | 37(±14) | 31(±12) | 22(±0) |
| $+O_2$ | $CuSO_4$ | 0.1 | - | 104(±5) | 119(±7) | 109(±3) | 108(±1) |
| $+O_2$ | $CuSO_4$ | 0.5 | - | 103(±2) | 116(±7) | 108(±5) | 106(±2) |
| $+O_2$ | $CuSO_4$ | 1 | - | 97(±1) | 110(±8) | 97(±3) | 96(±1) |
| $+O_2$ | $CuSO_4$ | 1.5 | - | 92(±4) | 87(±1) | 94(±5) | 95(±7) |
| $+O_2$ | $CuSO_4$ | 1.8 | - | 92(±3) | 86(±2) | 94(±7) | 96(±1) |
| $+O_2$ | $CuSO_4$ | 2 | - | 95(±5) | 90(±1) | 94(±4) | 94(±2) |
| $+O_2$ | $CuSO_4$ | 2.2 | - | 67(±2) | 64(±1) | 69(±6) | 84(±2) |
| $+O_2$ | $CuSO_4$ | 2.3 | - | 72(±1) | 66(±1) | 72(±3) | 83(±2) |
| $+O_2$ | $CuSO_4$ | 2.4 | - | 70(±1) | 64(±1) | 67(±2) | 74(±7) |
| $+O_2$ | $CuSO_4$ | 2.5 | - | 65(±2) | 67(±2) | 77(±1) | 90(±2) |
| $+O_2$ | $CuSO_4$ | 3 | - | 61(±2) | 54(±1) | 60(±4) | 67(±4) |
| $+O_2$ | $CuSO_4$ | 4 | - | 52(±4) | 42(±3) | 40(±1) | 39(±1) |
| $+O_2$ | $ZnSO_4$ | 0.05 | - | 97(±1) | 99(±2) | 100(±3) | 106(±2) |
| $+O_2$ | $ZnSO_4$ | 0.1 | - | 103(±2) | 103(±2) | 100(±3) | 105(±2) |
| $+O_2$ | $ZnSO_4$ | 0.5 | - | 97(±1) | 95(±3) | 96(±3) | 104(±2) |
| $+O_2$ | $ZnSO_4$ | 0.6 | - | 70(±2) | 79(±1) | 86(±4) | 92(±1) |
| $+O_2$ | $ZnSO_4$ | 0.7 | - | 55(±4) | 58(±7) | 72(±5) | 79(±1) |
| $+O_2$ | $ZnSO_4$ | 0.8 | - | 44(±3)[a] 48(±2)[b] | 56(±5)[a] 43(±2)[b] | 71(±1)[a] 57(±5)[b] | 82(±4)[a] 66(±3)[b] |
| $+O_2$ | $ZnSO_4$ | 1 | - | 23(±3) | 20(±2) | 30(±5) | 50(±3) |
| $+O_2$ | $ZnSO_4$ | 1.5 | - | 44(±5) | 14(±4) | 33(±37) | 15(±8) |
| $+O_2$ | EDTA | 0.1 | - | 81(±1) | 92(±3) | 91(±7) | 98(±1) |
| $+O_2$ | EDTA | 0.3 | - | 82(±2) | 87(±6) | 92(±3) | 91(±3) |
| $+O_2$ | EDTA | 0.5 | - | 88(±1) | 84(±2) | 84(±2) | 88(±1) |
| $+O_2$ | EDTA | 0.7 | - | 76(±1) | 69(±3) | 72(±2) | 73(±1) |
| $+O_2$ | EDTA | 1 | 81(±13) | 90(±7) | 89(±3) | 100(±9) | 79(±3) |
| $+O_2$ | EDTA | 1.3 | - | 73(±2) | 68(±3) | 70(±0) | 71(±1) |
| $+O_2$ | EDTA | 2 | - | 72(±2) | 68(±0) | 66(±2) | 70(±2) |
| $+O_2$ | $H_2O_2$ | 0.1 | - | 85(±5) | 87(±4) | 91(±2) | 90(±1) |
| $+O_2$ | $H_2O_2$ / $MnCl_2$ | 0.1 / 4 | - | 92(±5) | 94(±8) | 103(±6) | 87(±4) |
| $-O_2$ | EDTA | 1 | - | - | 70(±2) | - | - |
| $-O_2$ | DMG | 0.5 | - | - | 111(±2) | - | - |
| $-O_2$ | DMG | 1 | - | - | 104(±0.5) | - | - |
| $-O_2$ | $FeSO_4$ | 0.5 | - | - | 113(±4) | 111(±7) | - |
| $-O_2$ | $NiSO_4$ | 0.5 | - | - | 106(±2) | - | - |
| $-O_2$ | $NiSO_4$ | 1.5 | - | - | 54(±5) | - | - |
| $-O_2$ | $NiSO_4$ | 2 | - | - | 61(±14) | - | - |

**Supplementary Table 6**. Growth of *E. coli* strain JM109 (DE3) in response to various treatments. Mean growth of n = 3 biological replicates (± standard deviation) relative to mean of untreated cultures (n = 3 biological replicates) (for anaerobic 180 min treatment with 0.5 mM $FeSO_4$, n = 5 biological replicates for both treated and untreated cultures).

* One of the biological replicates in this experiment became contaminated, average of two biological replicates presented.

[a, b] This treatment performed with 3 biological replicates on two separate occasions.

| +/- $O_2$ | Treatment | Exposure time (min) | $\Delta C_{q\ nikA\text{-}rpoD}$ | *rpoD* $C_q$ difference relative to untreated aerobic samples <2? Yes/No | *rpoD* $C_q$ difference relative to corresponding (120 min) anaerobic untreated samples <2? Yes/No |
|---|---|---|---|---|---|
| $-O_2$ | 1 mM DMG | 120 | 0.27(±0.2) | Y | Y |
| $-O_2$ | 0.5 mM DMG | 120 | 0.38(±0.3) | Y | Y |
| $-O_2$ | 1 mM EDTA | 120 | 0.48(±0.2) | Y | Y |
| $-O_2$ | Untreated | 120 | 0.78(±0.1) | Y | n.a |
| $+O_2$ | 1 mM EDTA | 60 | 1.6(±0.5) | Y | n.a. |
| $+O_2$ | 1 mM EDTA | 120 | 2.1(±0.4) | Y | n.a. |
| $+O_2$ | Untreated | 120 | 2.3(±0.4) | n.a. | n.a. |
| $+O_2$ | 1 mM EDTA | 240 | 2.7(±0.5) | Y | n.a. |
| $+O_2$ | 0.5 mM EDTA | 120 | 3.4(±0.5) | Y | n.a. |
| $+O_2$ | 100 μM $H_2O_2$ | 120 | 3.6(±0.3) | Y | n.a. |
| $-O_2$ | 1.5 mM Ni | 120 | 4.5(±0.1) | Y | Y |
| $-O_2$ | 0.5 mM Ni | 120 | 5.1(±0.4) | Y | Y |
| $-O_2$ | 2 mM Ni | 120 | 5.9(±0.3) | Y | Y |
| $+O_2$ | 2 mM Ni | 120 | 6.7(±0.3) | N | n.a. |
| $+O_2$ | 3 mM Ni | 120 | 7.1(±0.3) | N | n.a. |
| $+O_2$ | 0.5 mM Ni | 120 | 7.2(±0.2) | Y | n.a. |

**Supplementary Table 7**. $\Delta C_q$ values obtained from qPCR analysis of indicated samples with primers specific to *nikA* and *rpoD* (control). $\Delta C_{q\ nikA\text{-}rpoD}$ obtained for each biological replicate, mean (± standard deviation). Data from samples with a difference in *rpoD* $C_q$ greater than 2 relative to untreated aerobic samples were not processed further.

| +/- $O_2$ | Treatment | Exposure time (min) | $\Delta C_{q\ mntS\text{-}rpoD}$ | *rpoD* $C_q$ difference relative to untreated aerobic samples <2? Yes/No |
|---|---|---|---|---|
| $+O_2$ | 1 mM EDTA | 60 | 0.16(±0.5) | Y |
| $+O_2$ | 1 mM EDTA | 240 | 0.66(±0.08) | Y |
| $+O_2$ | 1 mM EDTA | 120 | 0.82(±0.2) | Y |
| $+O_2$ | 0.5 mM EDTA | 120 | 0.92(±0.03) | Y |
| $+O_2$ | Untreated | 120 | 1.8(±0.7) | n.a. |
| $-O_2$ | Untreated | 120 | 2.3(±0.3) | Y |
| $+O_2$ | 1 mM EDTA | 20 | 3.3(±0.2) | N |
| $+O_2$ | 0.5 mM Mn | 120 | 3.9(±0.3) | Y |
| $+O_2$ | 2 mM Mn | 120 | 4.1(±0.3) | Y |
| $+O_2$ | 4 mM Mn | 120 | 4.3(±0.4) | Y |
| $+O_2$ | 100 µM $H_2O_2$ | 60 | 5.0(±0.3) | Y |
| $+O_2$ | 100 µM $H_2O_2$ | 120 | 5.2(±0.3) | Y |
| $+O_2$ | 100 µM $H_2O_2$ 4 mM Mn | 120 | 6.4(±0.7) | Y |
| $+O_2$ | 100 µM $H_2O_2$ 4 mM Mn | 60 | 6.9(±0.3) | Y |

**Supplementary Table 8**. $\Delta C_q$ values obtained from qPCR analysis of indicated samples with primers specific to *mntS* and *rpoD* (control). $\Delta C_{q\ mntS\text{-}rpoD}$ obtained for each biological replicate, mean (± standard deviation). Data from samples with a difference in *rpoD* $C_q$ greater than 2 relative to untreated aerobic samples were not processed further.

| +/- $O_2$ | Treatment | Exposure time (min) | $\Delta C_{q\ fepD\text{-}rpoD}$ | *rpoD* $C_q$ difference relative to untreated aerobic samples <2? Yes/No | *rpoD* $C_q$ difference relative to corresponding (120 or 180 min) anaerobic untreated samples <2? Yes/No |
|---|---|---|---|---|---|
| $+O_2$ | 1 mM EDTA | 60 | 1.8(±0.1) | Y | n.a. |
| $+O_2$ | 1 mM EDTA | 120 | 2.4(±0.6) | Y | n.a. |
| $+O_2$ | 0.5 mM EDTA | 120 | 2.7(±0.3) | Y | n.a. |
| $+O_2$ | 1 mM EDTA | 240 | 2.8(±0.1) | Y | n.a. |
| $+O_2$ | 1 mM EDTA | 20 | 3.2(±0.3) | N | n.a. |
| $-O_2$ | Untreated | 120 | 6.1(±0.4) | Y | n.a. |
| $-O_2$ | 0.5 mM Fe | 120 | 7.0(±0.04) | Y | Y |
| $+O_2$ | Untreated | 120 | 7.6(±0.16) | n.a. | n.a. |
| $+O_2$ | 0.5 mM Fe | 120 | 7.9(±0.1) | Y | n.a. |
| $+O_2$ | 100 μM $H_2O_2$ | 120 | 8.5(±0.4) | Y | n.a. |
| $-O_2$ | Untreated | 180 | 8.7(±0.3) | N | n.a |
| $-O_2$ | 0.5 mM Fe | 180 | 9.1(±0.3) | N | Y |

**Supplementary Table 9**. $\Delta C_q$ values obtained from qPCR analysis of indicated samples with primers specific to *fepD* and *rpoD* (control). $\Delta C_{q\ fepD\text{-}rpoD}$ obtained for each biological replicate, mean (± standard deviation). Data from samples with a difference in *rpoD* $C_q$ greater than 2 relative to untreated aerobic samples were not processed further, in samples treated with 0.5 mM $FeSO_4$ for 180 min or equivalent untreated samples this difference was not replicated with the use of an alternative control gene, *gyrA* (Supplementary Table 4, Methods).

| +/- $O_2$ | Treatment | Exposure time (min) | $\Delta C_{q\ rcnA\text{-}rpoD}$ | *rpoD* $C_q$ difference relative to untreated aerobic samples <2? Yes/No |
|---|---|---|---|---|
| $+O_2$ | 0.5 mM Co | 120 | 2.9(±0.4) | Y |
| $+O_2$ | 1 mM EDTA | 60 | 6.2(±0.2) | Y |
| $+O_2$ | 1 mM EDTA | 120 | 6.5(±0.2) | Y |
| $+O_2$ | 1 mM EDTA | 240 | 6.8(±0.3) | Y |
| $+O_2$ | 0.5 mM EDTA | 120 | 7.1(±0.2) | Y |
| $-O_2$ | Untreated | 120 | 7.9(±0.09) | Y |
| $+O_2$ | 1 mM EDTA | 20 | 7.9(±0.3) | N |
| $+O_2$ | Untreated | 120 | 8.1(±0.3) | n.a. |
| $+O_2$ | 100 µM $H_2O_2$ | 120 | 9.5(±0.5) | Y |

**Supplementary Table 10**. $\Delta C_q$ values obtained from qPCR analysis of indicated samples with primers specific to *rcnA* and *rpoD* (control). $\Delta C_{q\ rcnA\text{-}rpoD}$ obtained for each biological replicate, mean (± standard deviation). Data from samples with a difference in *rpoD* $C_q$ greater than 2 relative to untreated aerobic samples were not processed further.

| +/- $O_2$ | Treatment | Exposure time (min) | $\Delta C_{q\ znuA\text{-}rpoD}$ | *rpoD* $C_q$ difference relative to untreated aerobic samples <2? Yes/No |
|---|---|---|---|---|
| $+O_2$ | 1 mM EDTA | 60 | -2.1(±0.7) | Y |
| $+O_2$ | 1 mM EDTA | 240 | -2.0(±0.1) | Y |
| $+O_2$ | 1 mM EDTA | 120 | -1.9(±0.3) | Y |
| $+O_2$ | 1 mM EDTA | 20 | -1.8(±0.1) | N |
| $+O_2$ | 0.8 mM Zn | 120 | -1.7(±0.4) | Y |
| $+O_2$ | 0.5 mM EDTA | 120 | -1.4(±0.8) | Y |
| $+O_2$ | 0.6 mM Zn | 120 | 1.7(±0.1) | Y |
| $-O_2$ | Untreated | 120 | 2.1(±0.2) | Y |
| $+O_2$ | Untreated | 120 | 2.2(±0.3) | n.a. |
| $+O_2$ | 100 μM $H_2O_2$ | 120 | 4.3(±0.6) | Y |
| $+O_2$ | 1 mM Zn | 60 | 4.8(±0.2) | N |
| $+O_2$ | 1 mM Zn | 120 | 6.2(±0.2) | N |

**Supplementary Table 11**. $\Delta C_q$ values obtained from qPCR analysis of indicated samples with primers specific to *znuA* and *rpoD* (control). $\Delta C_{q\ znuA\text{-}rpoD}$ obtained for each biological replicate, mean (± standard deviation). Data from samples with a difference in *rpoD* $C_q$ greater than 2 relative to untreated aerobic samples were not processed further, in samples treated with 1 mM $ZnSO_4$ for 120 min this difference was not replicated with the use of an alternative control gene, *gyrA* (Supplementary Table 4, Methods).

| +/- $O_2$ | Treatment | Exposure time (min) | $\Delta C_{q\ zntA\text{-}rpoD}$ | *rpoD* $C_q$ difference relative to untreated aerobic samples <2? Yes/No |
|---|---|---|---|---|
| $+O_2$ | 2 mM Zn | 10 | -2.3(±0.1) | Y |
| $+O_2$ | 0.6 mM Zn | 120 | -1.9(±0.2) | Y |
| $+O_2$ | 1 mM Zn | 10 | -1.7(±0.3) | N |
| $+O_2$ | 1 mM Zn | 60 | -0.82(±0.1) | N |
| $+O_2$ | 1 mM Zn | 120 | -0.28(±0.3) | N |
| $+O_2$ | Untreated | 120 | 1.2(±0.4) | n.a. |
| $-O_2$ | Untreated | 120 | 1.7(±0.2) | Y |
| $+O_2$ | 0.8 mM Zn | 120 | 1.8(±0.3) | Y |
| $+O_2$ | 100 µM $H_2O_2$ | 120 | 2.7(±0.5) | Y |
| $+O_2$ | 1 mM EDTA | 60 | 5.8(±0.2) | Y |
| $+O_2$ | 1 mM EDTA | 120 | 6.5(±0.3) | Y |
| $+O_2$ | 0.5 mM EDTA | 120 | 6.6(±0.2) | Y |
| $+O_2$ | 1 mM EDTA | 20 | 7.0(±0.4) | N |
| $+O_2$ | 1 mM EDTA | 240 | 7.3(±0.2) | Y |

**Supplementary Table 12**. $\Delta C_q$ values obtained from qPCR analysis of indicated samples with primers specific to *zntA* and *rpoD* (control). $\Delta C_{q\ zntA\text{-}rpoD}$ obtained for each biological replicate, mean (± standard deviation). Data from samples with a difference in *rpoD* $C_q$ greater than 2 relative to untreated aerobic samples were not processed further, in samples treated with 1 mM $ZnSO_4$ for 120 min this difference was not replicated with the use of an alternative control gene, *gyrA* (Supplementary Table 4, Methods).

| +/- $O_2$ | Treatment | Exposure time (min) | $\Delta C_{q\ copA\text{-}rpoD}$ | *rpoD* $C_q$ difference relative to untreated aerobic samples <2? Yes/No |
|---|---|---|---|---|
| $+O_2$ | 2.4 mM Cu | 120 | -3.4(±0.07) | Y |
| $+O_2$ | 1.8 mM Cu | 120 | -2.7(±0.3) | Y |
| $-O_2$ | Untreated | 120 | -0.32(±0.2) | Y |
| $+O_2$ | Untreated | 120 | 1.3(±0.2) | n.a. |
| $+O_2$ | 1 mM EDTA | 60 | 1.4(±0.2) | Y |
| $+O_2$ | 1 mM EDTA | 120 | 1.6(±0.1) | Y |
| $+O_2$ | 0.5 mM EDTA | 120 | 1.8(±0.4) | Y |
| $+O_2$ | 1 mM EDTA | 240 | 2.2(±0.2) | Y |
| $+O_2$ | 100 µM $H_2O_2$ | 120 | 3.4(±0.3) | Y |
| $+O_2$ | 1 mM EDTA | 20 | 3.8(±0.4) | N |

**Supplementary Table 13**. $\Delta C_q$ values obtained from qPCR analysis of indicated samples with primers specific to *copA* and *rpoD* (control). $\Delta C_{q\ copA\text{-}rpoD}$ obtained for each biological replicate, mean (± standard deviation). Data from samples with a difference in *rpoD* $C_q$ greater than 2 relative to untreated aerobic samples were not processed further.

| +/- $O_2$ | Treatment | Exposure time (min) | $\Delta\Delta C_q$ |
|---|---|---|---|
| $+O_2$ | 1 mM EDTA | 60 | -6.7(±0.5) |
| $+O_2$ | 1 mM EDTA | 240 | -6.3(±0.08) |
| $+O_2$ | 1 mM EDTA | 120 | -6.1(±0.2) |
| $+O_2$ | 0.5 mM EDTA | 120 | -5.9(±0.2) |
| $+O_2$ | Untreated | 120 | -5.1(±0.7) |
| $-O_2$ | Untreated | 120 | -4.6(±0.3) |
| $+O_2$ | 0.5 mM Mn | 120 | -3.0(±0.3) |
| $+O_2$ | 2 mM Mn | 120 | -2.8(±0.3) |
| $+O_2$ | 4 mM Mn | 120 | -2.6(±0.4) |
| $+O_2$ | 100 µM $H_2O_2$ | 60 | -1.9(±0.3) |
| $+O_2$ | 100 µM $H_2O_2$ | 120 | -1.7(±0.3) |
| $+O_2$ | 100 µM $H_2O_2$, 4 mM Mn | 120 | -0.48(±0.7) |
| $+O_2$ | 100 µM $H_2O_2$, 4 mM Mn | 60 | - |

**Supplementary Table 14**. $\Delta\Delta C_q$ values for conditional response of *mntS* promoter. Determined by subtraction of $\Delta C_q$ for control condition (lowest expression, aerobic 60 min exposure to 100 µM $H_2O_2$ and 4 mM Mn, Supplementary Table 8) from $\Delta C_q$ for the condition of interest.

| +/- $O_2$ | Treatment | Exposure time (min) | $\Delta\Delta C_q$ |
|---|---|---|---|
| +$O_2$ | 1 mM EDTA | 60 | -6.7(±0.1) |
| +$O_2$ | 1 mM EDTA | 120 | -6.2(±0.6) |
| +$O_2$ | 0.5 mM EDTA | 120 | -5.9(±0.3) |
| +$O_2$ | 1 mM EDTA | 240 | -5.7(±0.1) |
| -$O_2$ | Untreated | 120 | -2.5(±0.4) |
| -$O_2$ | 0.5 mM Fe | 120 | -1.6(±0.04) |
| +$O_2$ | Untreated | 120 | -0.91(±0.2) |
| +$O_2$ | 0.5 mM Fe | 120 | -0.63(±0.1) |
| +$O_2$ | 100 μM $H_2O_2$ | 120 | - |

**Supplementary Table 15**. $\Delta\Delta C_q$ values for conditional response of *fepD* promoter. Determined by subtraction of $\Delta C_q$ for control condition (lowest expression, aerobic 120 min exposure to 100 μM $H_2O_2$, Supplementary Table 9) from $\Delta C_q$ for the condition of interest.

| +/- $O_2$ | Treatment | Exposure time (min) | $\Delta\Delta C_q$ |
|---|---|---|---|
| $+O_2$ | 0.5 mM Co | 120 | -6.6(±0.4) |
| $+O_2$ | 1 mM EDTA | 60 | -3.3(±0.2) |
| $+O_2$ | 1 mM EDTA | 120 | -3.0(±0.2) |
| $+O_2$ | 1 mM EDTA | 240 | -2.7(±0.3) |
| $+O_2$ | 0.5 mM EDTA | 120 | -2.4(±0.2) |
| $-O_2$ | Untreated | 120 | -1.6(±0.09) |
| $+O_2$ | Untreated | 120 | -1.4(±0.3) |
| $+O_2$ | 100 µM $H_2O_2$ | 120 | - |

**Supplementary Table 16**. $\Delta\Delta C_q$ values for conditional response of *rcnA* promoter. Determined by subtraction of $\Delta C_q$ for control condition (lowest expression, aerobic 120 min exposure to 100 µM $H_2O_2$, Supplementary Table 10) from $\Delta C_q$ for the condition of interest.

| +/- $O_2$ | Treatment | Exposure time (min) | $\Delta\Delta C_q$ |
|---|---|---|---|
| $+O_2$ | 1 mM EDTA | 60 | -5.6(±0.5) |
| $+O_2$ | 1 mM EDTA | 120 | -5.1(±0.4) |
| $+O_2$ | Untreated | 120 | -4.9(±0.4) |
| $+O_2$ | 1 mM EDTA | 240 | -4.4(±0.5) |
| $+O_2$ | 0.5 mM EDTA | 120 | -3.8(±0.5) |
| $+O_2$ | 100 μM $H_2O_2$ | 120 | -3.6(±0.3) |
| $+O_2$ | 0.5 mM Ni | 120 | - |

**Supplementary Table 17**. $\Delta\Delta C_q$ values for conditional response of *nikA* promoter under aerobic conditions. Determined by subtraction of $\Delta C_q$ for control condition (lowest expression, aerobic 120 min exposure to 0.5 mM Ni, Supplementary Table 7) from $\Delta C_q$ for the condition of interest.

| +/- $O_2$ | Treatment | Exposure time (min) | $\Delta\Delta C_q$ |
|---|---|---|---|
| $-O_2$ | 1 mM DMG | 120 | -5.7(±0.2) |
| $-O_2$ | 0.5 mM DMG | 120 | -5.6(±0.3) |
| $-O_2$ | 1 mM EDTA | 120 | -5.5(±0.2) |
| $-O_2$ | Untreated | 120 | -5.2(±0.1) |
| $-O_2$ | 1.5 mM Ni | 120 | -1.5(±0.1) |
| $-O_2$ | 0.5 mM Ni | 120 | -0.84(±0.4) |
| $-O_2$ | 2 mM Ni | 120 | - |

**Supplementary Table 18**. $\Delta\Delta C_q$ values for conditional response of *nikA* promoter under anaerobic conditions. Determined by subtraction of $\Delta C_q$ for control condition (lowest expression, anaerobic 120 min exposure to 2 mM Ni, Supplementary Table 7) from $\Delta C_q$ for the condition of interest.

| +/- $O_2$ | Treatment | Exposure time (min) | $\Delta\Delta C_q$ |
|---|---|---|---|
| +$O_2$ | 1 mM EDTA | 60 | -6.4(±0.7) |
| +$O_2$ | 1 mM EDTA | 240 | -6.3(±0.1) |
| +$O_2$ | 1 mM EDTA | 120 | -6.2(±0.3) |
| +$O_2$ | 0.8 mM Zn | 120 | -6.0(±0.4) |
| +$O_2$ | 0.5 mM EDTA | 120 | -5.7(±0.8) |
| +$O_2$ | 0.6 mM Zn | 120 | -2.6(±0.1) |
| -$O_2$ | Untreated | 120 | -2.2(±0.2) |
| +$O_2$ | Untreated | 120 | -2.1(±0.3) |
| +$O_2$ | 100 μM $H_2O_2$ | 120 | - |

**Supplementary Table 19**. $\Delta\Delta C_q$ values for conditional response of *znuA* promoter. Determined by subtraction of $\Delta C_q$ for control condition (lowest expression, aerobic 120 min exposure to 100 μM $H_2O_2$, Supplementary Table 11) from $\Delta C_q$ for the condition of interest.

| +/- $O_2$ | Treatment | Exposure time (min) | $\Delta\Delta C_q$ |
|---|---|---|---|
| $+O_2$ | 2 mM Zn | 10 | -9.6(±0.1) |
| $+O_2$ | 0.6 mM Zn | 120 | -9.1(±0.2) |
| $+O_2$ | Untreated | 120 | -6.1(±0.4) |
| $-O_2$ | Untreated | 120 | -5.6(±0.2) |
| $+O_2$ | 0.8 mM Zn | 120 | -5.5(±0.3) |
| $+O_2$ | 100 μM $H_2O_2$ | 120 | -4.5(±0.5) |
| $+O_2$ | 1 mM EDTA | 60 | -1.4(±0.2) |
| $+O_2$ | 1 mM EDTA | 120 | -0.7(±0.3) |
| $+O_2$ | 0.5 mM EDTA | 120 | -0.7(±0.2) |
| $+O_2$ | 1 mM EDTA | 240 | - |

**Supplementary Table 20**. $\Delta\Delta C_q$ values for conditional response of *zntA* promoter. Determined by subtraction of $\Delta C_q$ for control condition (lowest expression, aerobic 240 min exposure to 1 mM EDTA, Supplementary Table 12) from $\Delta C_q$ for the condition of interest.

| +/- $O_2$ | Treatment | Exposure time (min) | $\Delta\Delta C_q$ |
|---|---|---|---|
| $+O_2$ | 2.4 mM Cu | 120 | -6.8(±0.07) |
| $+O_2$ | 1.8 mM Cu | 120 | -6.0(±0.3) |
| $-O_2$ | Untreated | 120 | -3.7(±0.2) |
| $+O_2$ | Untreated | 120 | -2.1(±0.2) |
| $+O_2$ | 1 mM EDTA | 60 | -2.0(±0.2) |
| $+O_2$ | 1 mM EDTA | 120 | -1.8(±0.1) |
| $+O_2$ | 0.5 mM EDTA | 120 | -1.6(±0.4) |
| $+O_2$ | 1 mM EDTA | 240 | -1.2(±0.2) |
| $+O_2$ | 100 µM $H_2O_2$ | 120 | - |

**Supplementary Table 21**. $\Delta\Delta C_q$ values for conditional response of *copA* promoter. Determined by subtraction of $\Delta C_q$ for control condition (lowest expression, aerobic 120 min exposure to 100 µM $H_2O_2$, Supplementary Table 13) from $\Delta C_q$ for the condition of interest.

| Protein | Similarity (%) | Identity (%) |
|---|---|---|
| MntR | 93.0 | 89.8 |
| Fur | 97.3 | 96.7 |
| RcnR | 96.7 | 93.3 |
| NikR | 99.2 | 98.5 |
| Zur | 95.3 | 93.0 |
| ZntR | 97.9 | 92.2 |
| CueR | 94.2 | 89.1 |

**Supplementary Table 22**. Percentage similarity and identity between metal sensors of *E. coli* JM109 (DE3) and *Salmonella*. We do not note any variation for proposed metal binding residues between the *E. coli* and *Salmonella* proteins. Additionally, we note that residues required for allostery and DNA binding are conserved in MntR and Zur. Pairwise comparison performed with EMBOSS Needle (https://www.ebi.ac.uk/Tools/psa/emboss_needle/)[17].
